# Oncogenic PKA signaling stabilizes MYC oncoproteins via an aurora kinase A-dependent mechanism

**DOI:** 10.1101/2021.04.16.438110

**Authors:** Gary K. L. Chan, Samantha Maisel, Yeonjoo C. Hwang, Rebecca R. B. Wolber, Phuong Vu, Krushna C. Patra, Mehdi Bouhaddou, Heidi L. Kenerson, Raymond S. Yeung, Danielle L. Swaney, Nevan J. Krogan, Rigney E. Turnham, John D. Scott, Kimberly J. Riehle, Nabeel Bardeesy, John D. Gordan

**Affiliations:** Division of Hematology/Oncology, Helen Diller Family Comprehensive Cancer Center, University of California, San Francisco, San Francisco, CA, USA; Quantitative Biosciences Institute (QBI), University of California San Francisco, San Francisco, CA, USA; Department of Medicine, Harvard Medical School, Boston, Massachusetts, USA; Massachusetts General Hospital Cancer Center, Boston, Massachusetts, USA; Department of Cellular and Molecular Pharmacology, University of California San Francisco, San Francisco, CA, USA; David J. Gladstone Institute, San Francisco, CA, USA; Department of Surgery and Northwest Liver Research Program, University of Washington, Seattle, WA, USA; Department of Pharmacology, University of Washington Medical Center, Seattle, WA, USA

## Abstract

Genetic alterations that activate protein kinase A (PKA) signaling are found across many tumor types, but their downstream oncogenic mechanisms are poorly understood. We used global phosphoproteomics and kinome activity profiling to map the conserved signaling outputs driven by diverse genetic changes that activate PKA in human cancer, including melanoma and fibrolamellar carcinoma (FLC). We define two consensus networks of effectors downstream of PKA in cancer models. One is centered on RAS/MAPK components, and a second involves Aurora Kinase A (AURKA). We find that AURKA stabilizes c-MYC and n-MYC protein levels and expression programs in PKA-dependent tumor models, in part via a positive feedback loop mediated by the oncogenic kinase PIM2. This process can be suppressed by conformation-disrupting AURKA inhibitors such as CD-532. Our findings elucidate two independent mechanisms of PKA-driven tumor cell growth and offer insight into drug targets for study in FLC and other PKA-dependent malignancies.

## Introduction

Protein kinase A (PKA) is an evolutionarily conserved signaling effector with established roles in diverse physiological processes, including the regulation of growth, differentiation and metabolism (Turnham & Scott, 2016). Canonically PKA is controlled by cyclic AMP (cAMP) that are generated by the activation of the G protein-coupled receptor (GPCR) signaling. However genomic alterations in the components of the GPCR-PKA signaling pathway lead to constitutive activation of this kinase in many human diseases including cancer (Taylor et al., 2013). A variety of oncogenic events result in PKA stimulation, including ligand activation of upstream GPCRs (Coles et al., 2020; McCudden et al., 2005), point mutations in the G protein subunit *GNAS* (Iglesias-Bartolome et al., 2015; Patra et al., 2018), inactivation of PKA regulatory protein PKA-RIα (Yin et al., 2011), and activating mutations in *PRKACA*, which encodes the catalytic subunit of PKA (PKAc) itself (Berthon et al., 2015). Elevated PKA activity as a consequence of *GNAS* or *PRKACA* mutations has been reported in a variety of endocrine tumors (Salpea & Stratakis, 2014), including the *PRKACA L205R* mutation in adrenocortical and ACTH-producing pituitary tumors in patients with Cushing’s syndrome (Cao et al., 2014). Patients with germline inactivating mutations in *PRKAR1A* are predisposed to develop myxomas, thyroid and gonadal tumors, referred to as Carney Complex (Yin et al., 2011). Recently, a *DNAJB1-PRKACA* gene fusion has emerged as the dominant oncogenic event in a rare liver tumor, fibrolamellar carcinoma (FLC) (Honeyman et al., 2014). This gene fusion has been reported in 79-100% of FLC (Cornella et al., 2015; Honeyman et al., 2014), with rare cases instead bearing *PRKAR1A* deletion (Graham et al., 2018). *DNAJB1-PRKACA* fusions have also been described in very small subsets of hepatocellular carcinoma (HCC) (Cancer Genome Atlas Research Network., 2017), cholangiocarcinoma (Nakamura et al., 2015), and oncocytic biliary tumors (Singhi et al., 2020).

The PKA holoenzyme is composed of two catalytic (C) and two regulatory (R) subunits; its activation is regulated by the second messenger cAMP (Taylor et al., 2013). In its inactive state, R subunits form a homodimer that binds and inhibits the C subunits. cAMP generated by GPCR/GNAS-mediated stimulation of adenylyl cyclase binds R subunits, causing a conformational change that allows greater mobility and activity of the C subunits, while maintaining localization of active kinase complexes (Smith et al., 2017). This spatiotemporal specificity is provided by A-kinase anchoring proteins (AKAPs), which bind and sequester PKA to subcellular locations, creating microdomains for localized downstream signaling (Langeberg & Scott, 2015; Scott & Pawson, 2009).

PKA signaling is well established to modulate cancer-relevant processes including growth factor signaling, cell migration, cell cycle regulation and control of cell metabolism. However, it remains unclear which oncogenic pathways downstream of PKA are essential, and which tissues or contexts they have the greatest impact (Burton & McKnight, 2007; London et al., 2020). DNAJ-PKAc stimulates ERK activation in an FLC model system (Turnham et al., 2019), operating via its interaction with AKAP-Lbc (Smith et al., 2010). In *GNAS*-mutant pancreatic tumor cells, PKAc-mediated suppression of the salt-inducible kinases (SIK1-3) supports tumor growth (Patra et al., 2018). PKAc has also been connected to control of G2/M transition (Grieco et al., 1996; Kotani et al., 1998) and to cell survival under glucose starvation (Palorini et al., 2016). Interestingly, PKA also has clear, but context-specific, tumor suppressive functions involving modulation of the Hedgehog and Hippo signaling pathways, and is mutationally inactivated in a subset of cancers (Iglesias-Bartolome et al., 2015; Tokita et al., 2019).

Despite its oncogenic action in multiple tumor types, PKAc is challenging to target directly. Consistent with the critical, widespread role of PKAc in normal physiology, mice with *PRKACA* deletion have less than 30% survival into adulthood (Skålhegg et al., 2002). Thus, a better understanding of the essential PKA effectors in individual tumor types is potentially a more tractable path for therapeutic development. In order to gain insight into oncogenic PKA signaling networks and to identify pharmacologically targetable downstream effects, we have explored its downstream kinases. We generated cell models with regulatable PKA activity and used them to produce a proteomic profile of PKA signaling. We show that elevated PKA signaling in models with genetic activation of the pathway, including FLC, promotes cell proliferation by increasing expression of c-MYC and n-MYC. In addition, we identify and validate a class of Aurora Kinase A (AURKA) inhibitors capable of blocking this process and PKA-driven tumor growth.

## Results

### *PRKACA* alterations are common among cancer types

We first sought to characterize the frequency and pattern of somatic mutations that impact PKA activity in human cancer in addition to FLC, which carries a unique *DNAJB1-PRKACA* fusion transcript in 79-100% of cases (Cornella et al., 2015; Honeyman et al., 2014). We analyzed the frequency of PKA-activating somatic alterations in different types of human cancer reported by TCGA Pan Cancer Project (Weinstein et al., 2013), including both *PRKACA* gain-of-function and *PRKAR1A* loss-of-function mutations in addition to copy number alterations (Fig. 1A). We found an incidence of *PRKACA* amplification of 0.3-3.2%, and a rate of activating mutations of 0.2-2.7% across multiple cancers. The greatest frequency of activation occurred in malignant peripheral nerve sheath tumors (MPNST) and ovarian cancers at 11.1 and 11.3%, respectively. *PRKAR1A* loss of function was rarer, including both inactivating mutations (0.2-0.7%) and deep deletions (0.4-0.6%) that were predominantly detected in adrenocortical carcinoma (5.3% and 4.0%, respectively; Fig. 1B).

**Figure 1.**
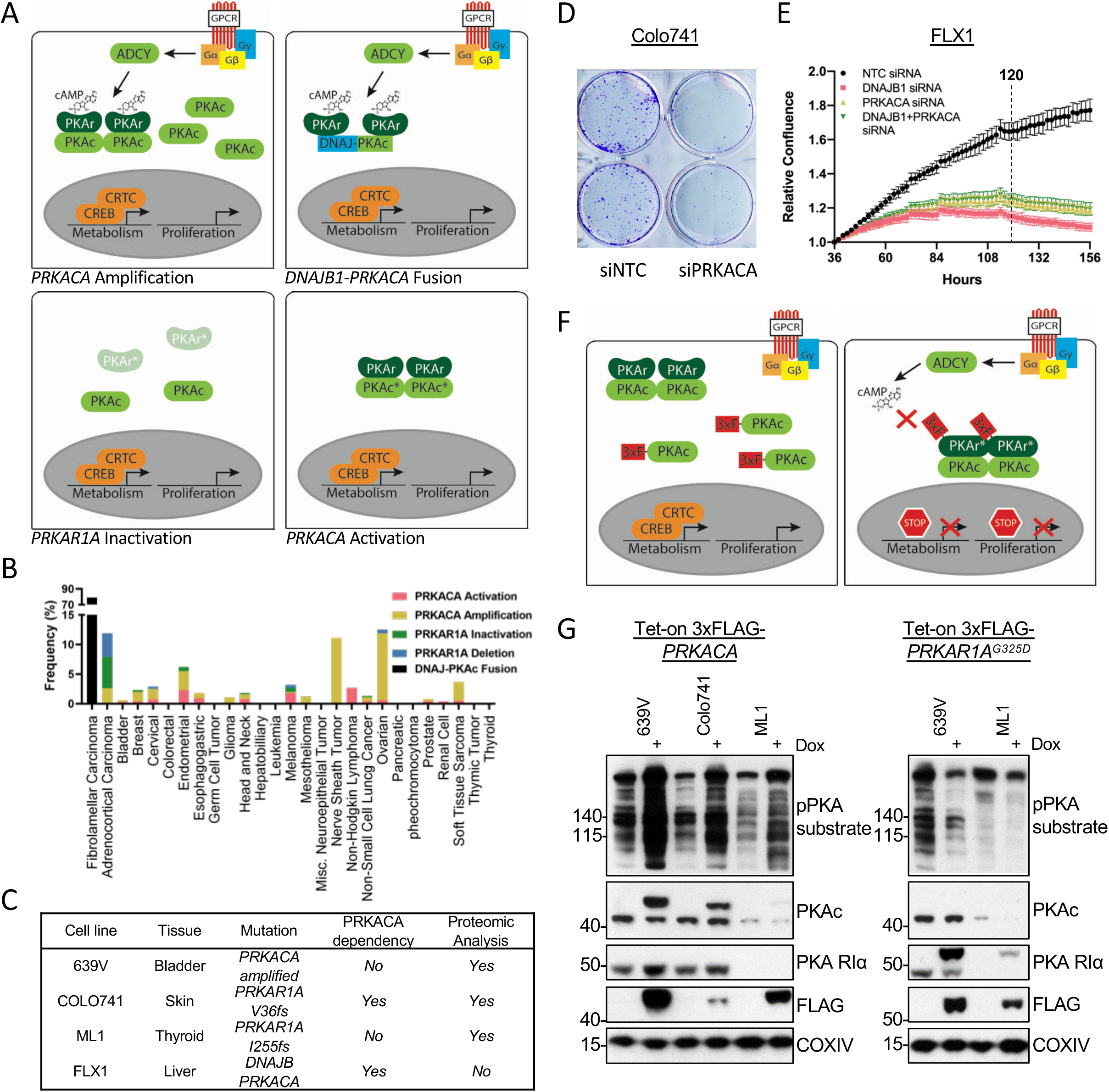
Recurrent PKA activating somatic alterations in human cancer. (A) Pathway illustrations of different PKA activating genomic alterations. Top left: *PRKACA* amplification. Top right: *DNAJB1-PRKACA* fusion. Bottom left: *PRKAR1A* inactivation or deletion. Bottom right: *PRKACA* activation. (B) TCGA PanCancer Project analysis shows the frequency of *PRKACA* gain-of-function (red and yellow) and *PRKAR1A* loss-of-function (green and blue) alterations in various cancer types. The reported frequency of *DNAJB1-PRKACA* fusion in FLC clinical samples is also included. (C) Cell lines used in this study, their PKA related mutation, *PRKACA* dependency and inclusion in proteomic analyses. (D) Clonogenic assay of Colo741 cells treated with *NTC* (left) or *PRKACA* (right) siRNA. Experiments were done twice with different cell density, and two of the three technical replications from the best representation were shown. (E) Relative confluence of FLX1 cells in 96 wells plate treated with *NTC, DNAJB1, PRKACA* or a mixture (50:50) of *DNAJB1* and *PRKACA* siRNA. 120 hours is marked with the dashed line. Result was the mean ±s.d of one biological replicate, n = 15 for each condition. (F) Schematic of doxycycline-induced PKAc modulating system. Left: Tet-on 3xFLAG-*PRKACA*. Right: Tet-on 3xFLAG-*PRKAR1A*^*G325D*^. (G) Immunoblots showing the change of PKA activity, as indicated by phospho-PKA substrate, in different cell lines with dox-inducible 3xFLAG-*PRKACA* or *PRKAR1A*^*G325D*^ with 1 ug/ml doxycycline (dox) for 48 hours. Left: transgenic cell lines with Tet-on 3xFLAG-*PRKACA*. Right: transgenic cell lines with Tet-on 3xFLAG-*PRKAR1A*^*G325D*^.

We next obtained cell models with PKA-activating mutations to study the signaling effects of PKA through engineered PKAc activation or inhibition, as these cells should have intact oncogenic signaling downstream of PKA (Fig. 1C). These models include 639V (bladder cancer with *PRKACA* amplification (Barretina et al., 2012)) and Colo741 and ML1 (skin and thyroid cancer models with *PRKAR1A* frameshift mutations (Ghandi et al., 2019)). Each of these models has been profiled for PKA dependency in the context of the Cancer Dependency Map program, with Colo741 specifically being highly dependency on PRKACA (data accessed from depmap.org) (McFarland et al., 2018). We confirmed the dependency of Colo741 cells on PKAc by clonogenic assay following treatment with *PRKACA* siRNA or *NTC* (non-targeting control; Fig. 1D). We also test PKAc dependency in a novel cell line from a patient-derived xenograft model of FLC, designated FLX1 cells (Oikawa et al., 2015). FLX1 cells were treated with siRNA for *DNAJB1, PRKACA*, an equal part mixture of *DNAJB1* and *PRKACA*, or *NTC* for 36 hours, and then quantified by real-time microscopy. siRNA targeting either or both components of the fusion reduced cell proliferation rate by 63 to 79% compared to *NTC* at 120 hours, demonstrating the dependency of these cells on the *DNAJB1-PRKACA* fusion (Fig. 1E).

### Kinome profile of oncogenic PKA signaling

To create stably inducible cell models for proteomic analysis, we introduced doxycycline (dox)-controlled 3xFLAG*-PRKACA* or *PRKAR1A*^*G325D*^ into 639V, Colo741, and ML1 cells. Expression of PKA subunits was via lentiviral infection. The *PRKAR1A*^*G325D*^ mutant exhibits impaired binding to cAMP, and thus serves as a dominant negative inhibitor of PKAc (Viste et al., 2005; Willis et al., 2011) (Fig. 1F). We confirmed that these constructs modulated PKA activity following dox induction, as assessed by immunoblot analysis using a phospho-PKA substrate antibody (Fig. 1G). We were not able to maintain the *PRKAR1A*^*G325D*^ transgene in the Colo741 cell line, possibly due to leaky expression resulting in reduced cell viability via chronic partial inhibition of PKAc.

We used a combination of proteomic strategies and computational integration to screen for oncogenic kinases downstream of PKA. The engineered cells described above were cultured with or without dox for 48 hours and analyzed with our global phosphoproteomics and Multiplex Inhibitor Bead (MIBs) pipelines (Budzik et al., 2020; Coles et al., 2020; Donnella et al., 2018; Sos et al., 2014). Further bioinformatic analysis was performed on the global phosphoproteomic data set with the Phosfate analysis tool to infer changes in kinase activity (Ochoa et al., 2016). These complementary tools allow us to measure known kinase/substrate relationships (Phosfate) and directly assay the activity of kinases whose substrates are not well known (MIBs). We initially confirmed the desired impact of *PRKACA* and *PRKAR1*^*G325D*^ transgenes: using the 639V transgenic cell lines, we showed that the phosphorylation levels of the PKAc target SIK2 pS587 (Patra et al., 2018) increased with PKAc induction and decreased with PKA R1α^G325D^ induction in our global phosphoproteomics analysis (Fig. 2A). Similarly, we detected upregulation of PKAc in both Phosfate and MIBs results following dox treatment of PKAc-inducible cells (Fig. 2B).

**Figure 2.**
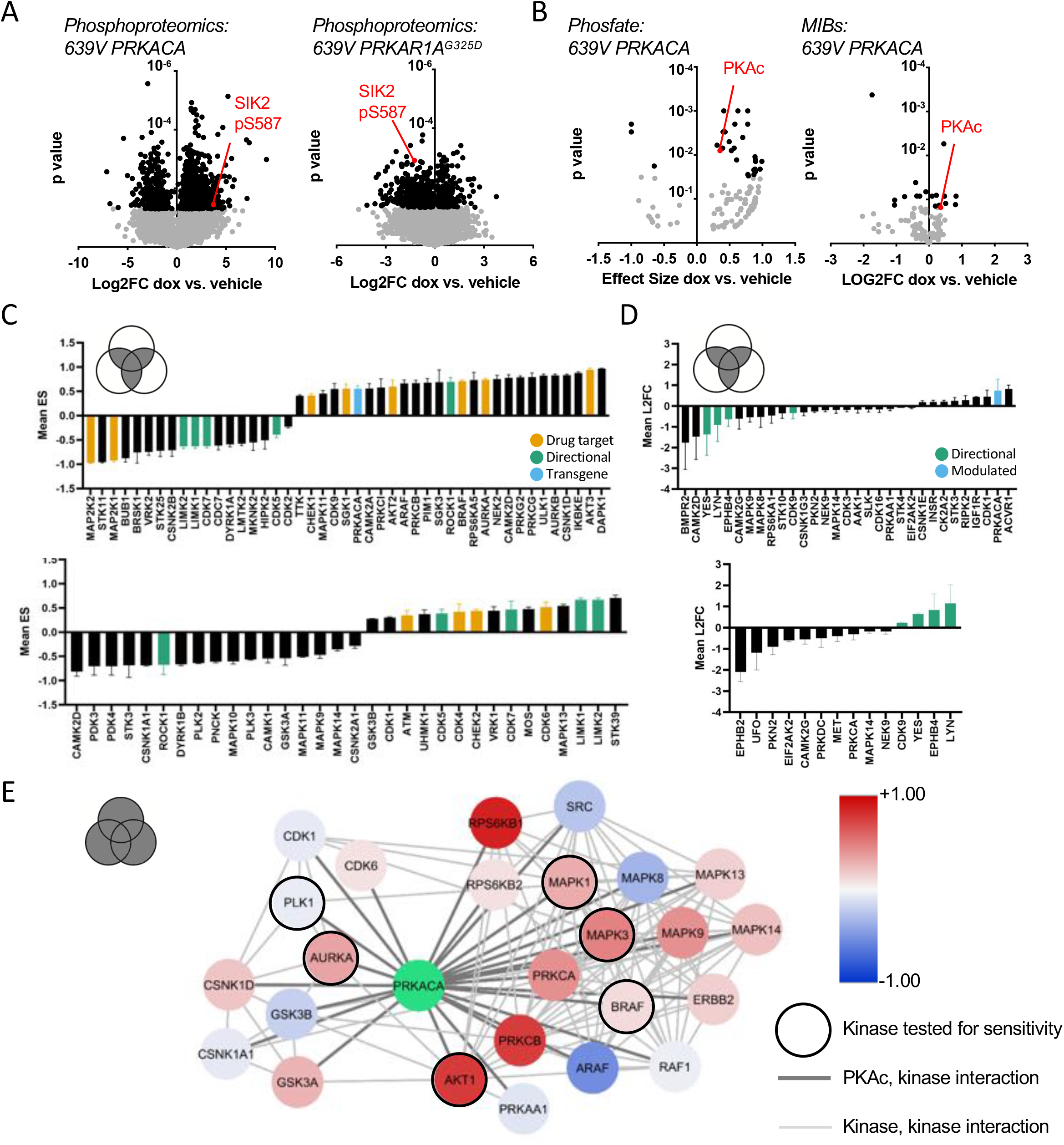
Kinome profiling to identify signaling nodes downstream of *PRKACA*. (A) Global phosphorylation change based on combined analysis of Phosfate and MIBs using 639V with *PRKACA* or *PRKAR1A*^*G325D*^ induced. (B) Change in kinase activity based on Phosfate analysis or MIBs pipeline using 639V Tet-on 3xFLAG-*PRKACA* compared to control. (C) In intersect, inferred kinase activity changes based on global phosphoproteomics using the Phosfate analysis tool averaged across different transgenic cell lines, with versus without dox induction. (top) 639V, Colo741, and ML1 cells with Tet-on 3xFLAG-*PRKACA*. (Bottom) 639V and ML1 with Tet-on 3xFLAG-*PRKAR1A*^*G325D*^. (D) In intersect, kinase activity changes based on MIBS averaged across different transgenic cell lines, with versus without dox induction. (top) 639V, Colo741, and ML1 cells with Tet-on 3xFLAG-*PRKACA*. (Bottom) 639V and ML1 with Tet-on 3xFLAG-*PRKAR1A*^*G325D*^. (E) MIBs and Phosfate kinome profiles from 639V, Colo741 and ML1 with Tet-on 3xFLAG-*PRKACA* and 639V and ML1 with Tet-on 3xFLAG-*PRKAR1A*^*G325D*^ were integrated by network propagation. Each node was filled to represent the original z-scores of kinase activity, which were averaged across cell lines.

We next integrated the proteomics data in 4 categories: Phosfate for cells with inducible PKAc (Fig. 2C, top), Phosfate for cells with inducible PKA R1α^G325D^ (Fig. 2C, bottom), MIBs for cells with inducible PKAc (Fig. 2D, top), and MIBs for cells with inducible PKA R1α^G325D^ (Fig. 2D, bottom). Across these datasets, YES, LYN, EPHB4, LIMK1, LIMK2, CDK5, and CDK7 activity was reduced following PKAc overexpression and increased following PKAc inhibition by PKA R1α ^G325D^ expression; conversely, we saw that ROCK1 was upregulated following PKAc overexpression and downregulated following PKA R1α^G325D^ expression. These results provide important proof of concept for our genetic model system and are consistent with extensive data connecting PKAc and cell migration (Howe, 2004). Since agents targeting these kinases have yet to find application in cancer treatment, we focused credentialed drug targets among the list of candidates, based on their potential near-term therapeutic significance. In this regard, we observed upregulation of multiple pro-proliferative kinases including AURKA, BRAF, and AKT2 (marked in yellow); unexpectedly MAP2K2 was downregulated. Interestingly, we also observed downregulation of the tumor suppressor STK11 by PKAc. Fewer signaling changes influencing proliferation resulted from *PRKAR1A*^*G325D*^ induction (Fig. 2C bottom, 2D bottom), although we note that our cell models have other oncogenic mutations acting on PKAc-regulated pathways that could maintain their activity despite PKA R1α^G325D^ expression.

Differences in isoform expression and redundant functions of kinases can obscure relationships between proteomic data sets when they are combined as above. Thus, we also used network propagation to integrate data across all of our cell models and both proteomic platforms (Cowen et al., 2017), using established pathway relationships from the ReactomeFI network to define functional connections between activate kinases within the PKA-regulated kinome (Wu & Haw, 2017). We then imported the propagated network into Cytoscape to visualize PKAc and its direct neighbors that are significantly altered when *PRKACA* or *PRKAR1A*^*G325D*^ was induced (Fig. 2E). We formatted the analysis to show PKAc effects by altering the directionality of *PRKAR1A*^*G325D*^, such that kinases that are upregulated by PKAc function in either data set are marked as positive (red), and downregulated negative (blue). This analysis defined two functional PKA-dependent clusters with both networks including potential drug targets (marked with black borders). One is marked by growth factor signaling effectors such as BRAF, multiple MAPKs, AKT, PKCs as well as the receptor kinase ERBB2. Interestingly, a second network emerged, notable for multiple cell cycle kinases involved in the regulation of G2/M including AURKA, PLK1, GSK3α/β, and several casein kinase family members.

We next assessed alterations in how PKAc overexpression affects sensitivity to inhibitors of these kinases in our ML1 cell model (Fig. S1A) There were three key findings: First, our analysis showed a minor decrease in sensitivity to MAPK pathway-targeted agents with PKAc induction. Second, we detected little impact on DNA-damage response targets. Third, we noted that PKAc induction increased sensitivity to the newly developed conformation-disrupting AURKA inhibitors (CD-AURKAi) CD-532 and MLN-8237. In contrast to the latter pharmacological effects, PKAc induction was unaffected by the older generation clinical candidate ENMD-2076 (Fig. S1B).

### AURKA inhibition with CD532 modulates c-MYC and n-MYC expression and cell proliferation

We next screened this panel of AURKA inhibitors against the PKAc-dependent Colo741 and FLX1 cell lines. CD-AURKAi CD532 had a more marked effect on cell viability than any other compound (Fig. 3A) (Dauch et al., 2016; Gustafson et al., 2014). Importantly, drug sensitivities (EC50 = 217.3 nM for Colo741; 692.8 nM for FLX1) matched the reported dose range for inhibition of AURKA kinase activity (146.7-223.2 nM) (Gustafson et al., 2014). CD-AURKAi have been linked to MYC protein degradation by displacing MYC from AURKA containing complexes and allowing its ubiquitylation (Dauch et al., 2016; Gustafson et al., 2014). Accordingly, CD532 or MLN8237 treatment nearly abolished c-MYC expression in Colo741 cells; CD532 was more active than MLN8237 in FLX1 in reducing by n-MYC and c-MYC expression (Fig. 3B). qRT-PCR showed that treatment with CD532 for 24hrs did not affect *MYC or MYCN* mRNA levels, consistent with an effect on stability of c-MYC and n-MYC proteins. In contrast, we observed significant downregulation of MYC target genes, including *E2F1* (p value = 0.0279) and *VCAN* (p value = 0.0491) (Fig. 3C).

**Figure 3.**
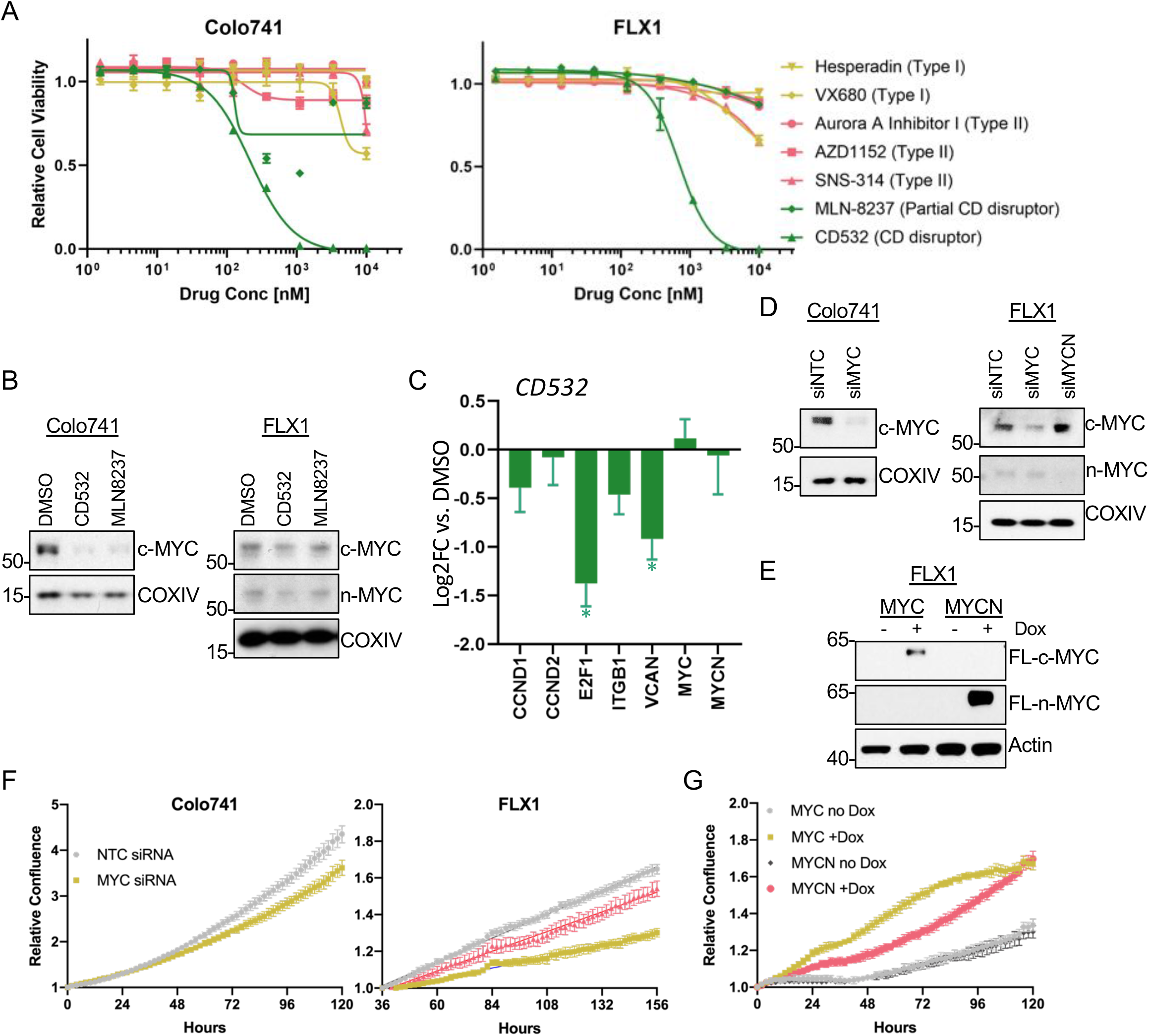
c-MYC and n-MYC promote proliferation of PKA-dependent cell models. (A) Colo741 and FLX1 cells in 96-well plates were treated with various types of AURKA inhibitors for 72h at different concentrations, and their relative cell viability was measured by CTG assay versus untreated control samples. Results are the mean ±s.e.m of triple biological replicates, three technical replicates per biological replicate. Inhibitors are colored based on their binding mode. (B) Immunoblots showing the expression of c-MYC and n-MYC in Colo741 and FLX1 cells after treating with CD532 or MLN8237 for 24 hours. (C) RT-qPCR analysis of *MYC, MYCN* and their downstream genes in FLX1 cells treated with 1uM CD532 for 24 hours. Results are the mean ±s.e.m of three biological replicates, three technical replicates for each biological replicate, targets with P value <0.05 form paired t-test for CD532 treated versus DMSO were labelled with the asterisk. (D) Immunoblots showing the change of c-MYC and n-MYC levels in Colo741 and FLX1 cells after *MYC or MYCN* siRNA knockdown for 48 hours. (E) Immunoblots showing the induction of Tet-on 3xFLAG-*MYC* and *MYCN* in respective FLX1 transgenic cells after dox induction for 48 hours. (F) Relative confluence of Colo741 and FLX1 cells in 96 well plates after *MYC* or *MYCN* knockdown with siRNA. FLX1 cells were incubated 36 hours before recording to ensure better adhesion. Experiments were done in duplicate and representative results were shown with mean ±s.d, n = 10 for each condition. (G) Relative confluence of transgenic FLX1 cells with doxycycline-controlled Tet-on 3xFLAG-*MYC* or *MYCN* in 96 well plates after treatment with or without 1ug/ml dox. Confluence was imaged for 120 hours and analyzed by Incucyte. Experiment was duplicate, the representative results shown with mean ±s.d, n = 6 for each condition.

Consistent with the functional relevance of MYC protein regulation downstream of PKAc, siRNA-mediated knockdown of c-MYC and n-MYC suppressed proliferation in Colo741 cells. Similar results were observed upon c-MYC siRNA knockdown in FLX1 cells (n-MYC is not expressed in Colo741; Fig. 3D, F). Conversely, ectopic expression of 3xFLAG-*MYC* or 3xFLAG-*MYCN* using a doxycycline-controlled system increased proliferation of FLX1 cells (Fig. 3E, G). Thus, MYC-family proteins are important regulators of proliferation in PKA-dependent cancers.

### *PRKACA* mediates c-MYC and n-MYC expression

Although the role of AURKA in maintaining c-MYC and n-MYC stability has been previously reported (Dauch et al., 2016; Gustafson et al., 2014; Otto et al., 2009), it has not been directly connected to upstream oncogenic signaling. Accordingly, we examined whether PKAc induces c-MYC or n-MYC expression in PKA-driven cancer cell lines using gain-and loss-of-function approaches. Based on their dependence on PKA for proliferation, distinct PKA activating mutations and similar stabilization of MYC family members in response to PKA activation, we again used the Colo741 and FLX1 cell lines for these analyses.

To hyperactivate PKAc, we treated cells with forskolin (FSK), an adenylyl cyclase stimulator, and IBMX, a phosphodiesterase inhibitor. This drug combination generates sustained and supraphysiological production of intracellular cAMP. Immunoblot of cells treated for 0.5, 2, or 4 hours revealed rapid activation of PKA that correlated with progressive c-MYC and n-MYC stabilization (Fig. 4A) although AURKA levels were unchanged (Fig. S2). Conversely, siRNA targeting *PRKACA* in Colo741 and FLX1 resulted in attenuated PKA substrate phosphorylation and progressive decreases in c-MYC and n-MYC (when expressed) (Fig. 4B). Additionally, we generated an FLX1 cell line with the dox-inducible 3xFLAG-*PRKAR1A*^*G325D*^ transgene, in which produced the expected reductions in PKA substrate phosphorylation and c-MYC and n-MYC levels (Fig. 4C).

**Figure 4.**
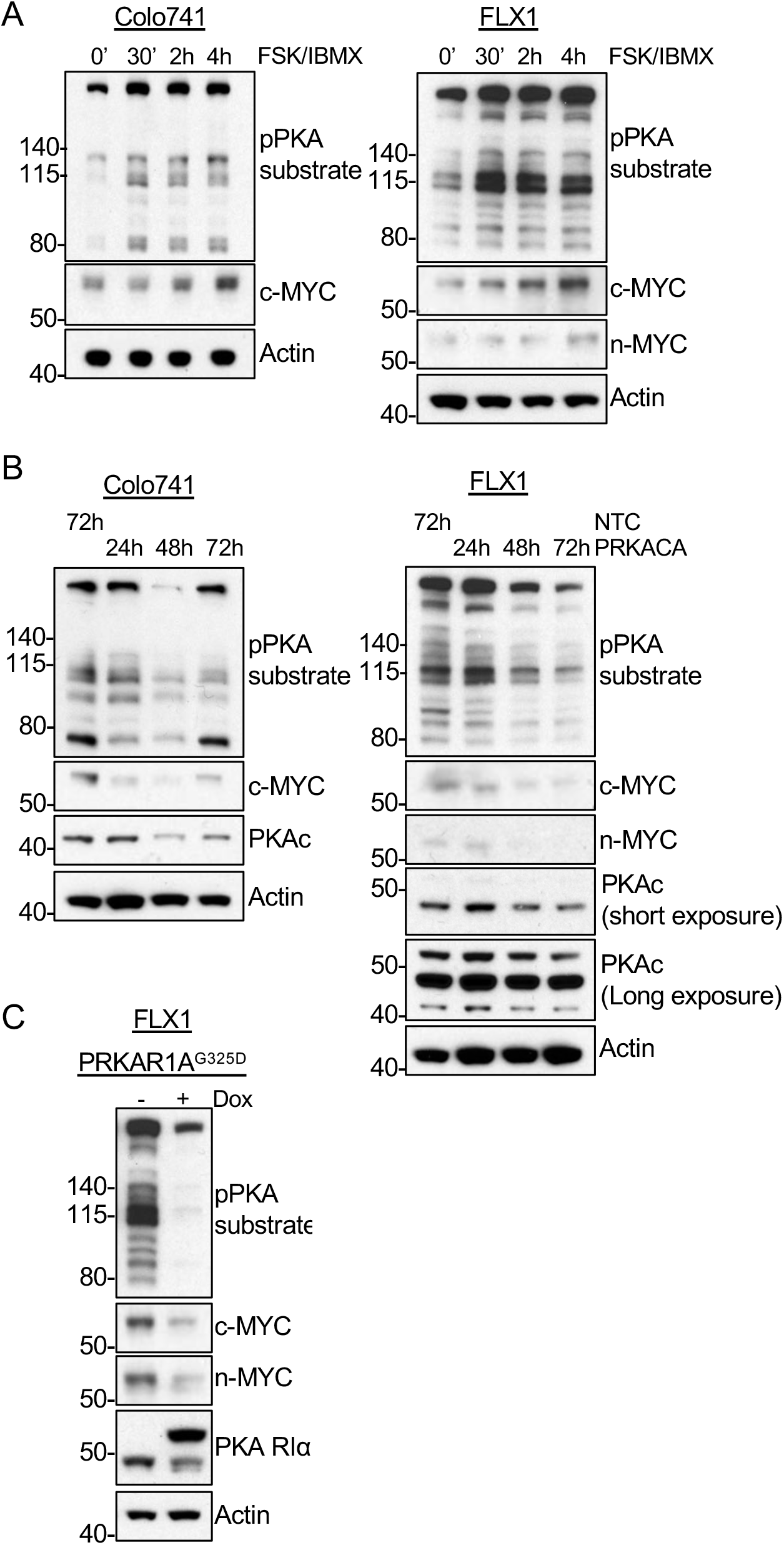
PKA activity increases c-MYC and n-MYC protein levels. (A) Immunoblots showing the change of PKA activity, as indicated by phospho-PKA substrate, and c-MYC and n-MYC expression in Colo741 and FLX1 cells after treating with 50uM FSK and 50uM IBMX for 0, 30 minutes, 2 and 4 hours. (B) Immunoblots showing the change of PKA activity and PKAc, c-MYC and n-MYC expressions in Colo741 and FLX1 cells after *PRKACA* siRNA knockdown for 24, 48 and 72 hours vs. 72 hours with non-targeting control (NTC). (C) Immunoblots showing the change of PKA activity and c-MYC and n-MYC levels in transgenic FLX1 cells with Tet-on 3xFLAG-*PRKAR1A*^*G325D*^ with or without dox for 72 hours.

### PIM2 also regulates c-MYC and n-MYC expression in PKA-driven cells

We further investigated the possibility that other kinases might act synergistically with AURKA to regulate MYC stability downstream of PKA. Using FLX1 cells, we screened a kinome-wide siRNA library for modifiers of cellular proliferation and identified 30 kinases whose genetic depletion reduced cell proliferation with a Z score < -1, and 20 kinases that increased proliferation (Fig. 5A). As indicated in Fig. 5A, we focused on non-metabolic kinases that reduced cellular proliferation and chose the most prominent candidate, *PIM2*, to directly test its effects on c-MYC expression in FLX1. PIM2 is a serine/threonine kinase with similar substrates and function to AKT, and is also known to play a role in MYC regulation (Fox et al., 2003; Hammerman et al., 2005; Zhang et al., 2008). The related kinase PIM1 was identified by the Phosfate analytical tool as an enzyme whose activity that is elevated in response to PKAc overexpression (Fig. 2C). This led to our working hypothesis that PKA-dependent mobilization of PIM kinases contributes to MYC expression. Consistent with this notion, we found that PIM2 knockdown specifically reduced c-MYC expression in Colo741 cells, while PIM1 did so in FLX1 cells (Fig. 5B). Next, we investigated a functional interplay between PIM kinases and AURKA inhibition. We found the PIM inhibitor CX-6258 alone reduced expression of c-MYC and n-MYC in both cell systems, and had a cooperative effect with MLN8237 in FLX1 cells (Fig. 5C).

**Figure 5.**
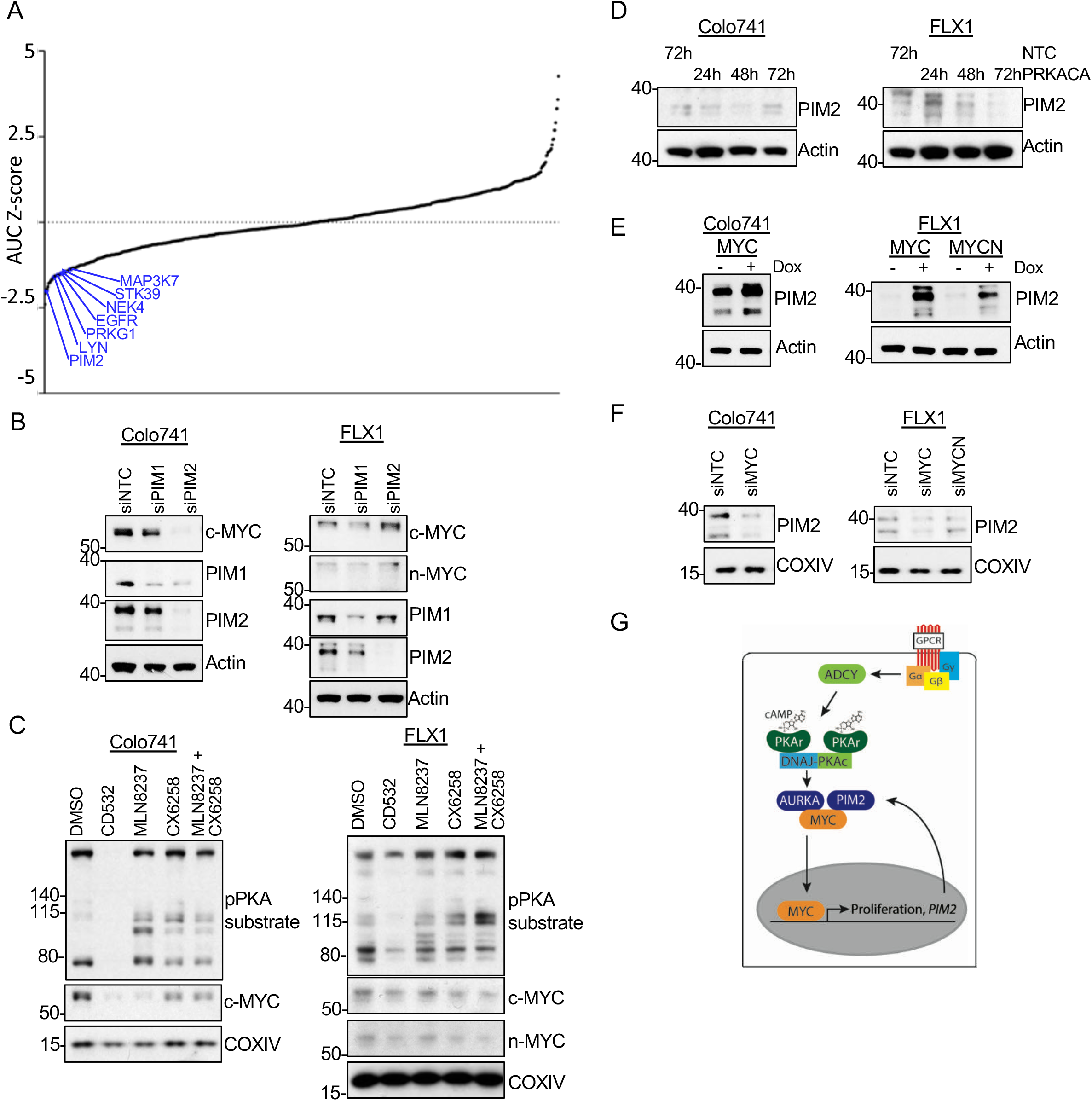
PIM2 increases c-MYC /n-MYC expression in cooperation with AURKA. (A) siRNA kinase library screen with FLX1 in 384 well plates for 7 days shows the effect of each target kinase on cell proliferation (average of 3 biological replicates). Selected non-metabolic kinases that decrease cell proliferation with z-score < -1 were marked. (B) Immunoblots showing the change of c-MYC and n-MYC levels in Colo741 and FLX1 cells after *PIM1* or *PIM2* siRNA knockdown for 48 hours. (C) Immunoblot showing the change of PKA activity, as indicated by phospho-PKA substrate, and c-MYC and n-MYC levels in Colo741 and FLX1 cells after treating with DMSO, 1uM CD532, 1uM MLN8237, 1uM CX6258 or combination of 1uM MLN8237 and 1uM CX6258 for 24 hours. (D) Immunoblots showing the change of PIM2 expression in Colo741 and FLX1 cells after *PRKACA* siRNA knockdown for 24, 48 and 72 hours and NTC for 72 hours. (E) Immunoblots showing the change of PIM2 level after *MYC* or *MYCN* overexpression in Colo741 or FLX1 transgenic cells after dox induction for 48 hours. (F) Immunoblots showing the change of PIM2 levels in Colo741 and FLX1 cells after *MYC* or *MYCN* siRNA knockdown for 48 hours. (G) Schematic of DNAJ-PKAc mediating cell proliferation in FLC by stabilizing MYC family proteins via AURKA and producing a positive feedback loop with PIM2.

To determine whether PIM2 is also responsive to PKA activation, Colo741 and FLX1 cells were treated with *PRKACA* siRNA or *NTC* for 24, 48 and 72 hours. PIM2 levels were reduced in Colo741 and FLX1 when *PRKACA* was knocked down (Fig. 5D). Moreover, ectopic expression of 3xFLAG*-MYC* or 3xFLAG*-MYCN* provoked a significant increase in PIM2 in these cells (Fig. 5E). Conversely, siRNA mediated depletion of *MYC* reduced PIM2 levels in both Colo741 and FLX1 cells (Fig. 5F). MYCN siRNA gene silencing in FLX1 cells had little effect (Fig. 5F). These results support a model where PIM2 is responsive to PKA activity via its effect on c-MYC and n-MYC, and then feeds back to stabilize MYC (Fig. 5G).

### PKA/MYC relationship in isogenic FLC models and human FLC samples

To confirm the relationship between PKA and MYC in other FLC models, we used the recently described isogenic FLC model (Turnham et al., 2019), in which the *DNAJB1-PRKACA* fusion was CRISPR engineered into AML12 immortalized murine hepatocytes. We once again saw that the engineered FLC clone had increased basal PKA activation, as well as higher c-MYC, n-MYC and PIM2 expression (Fig. 6A).

**Figure 6.**
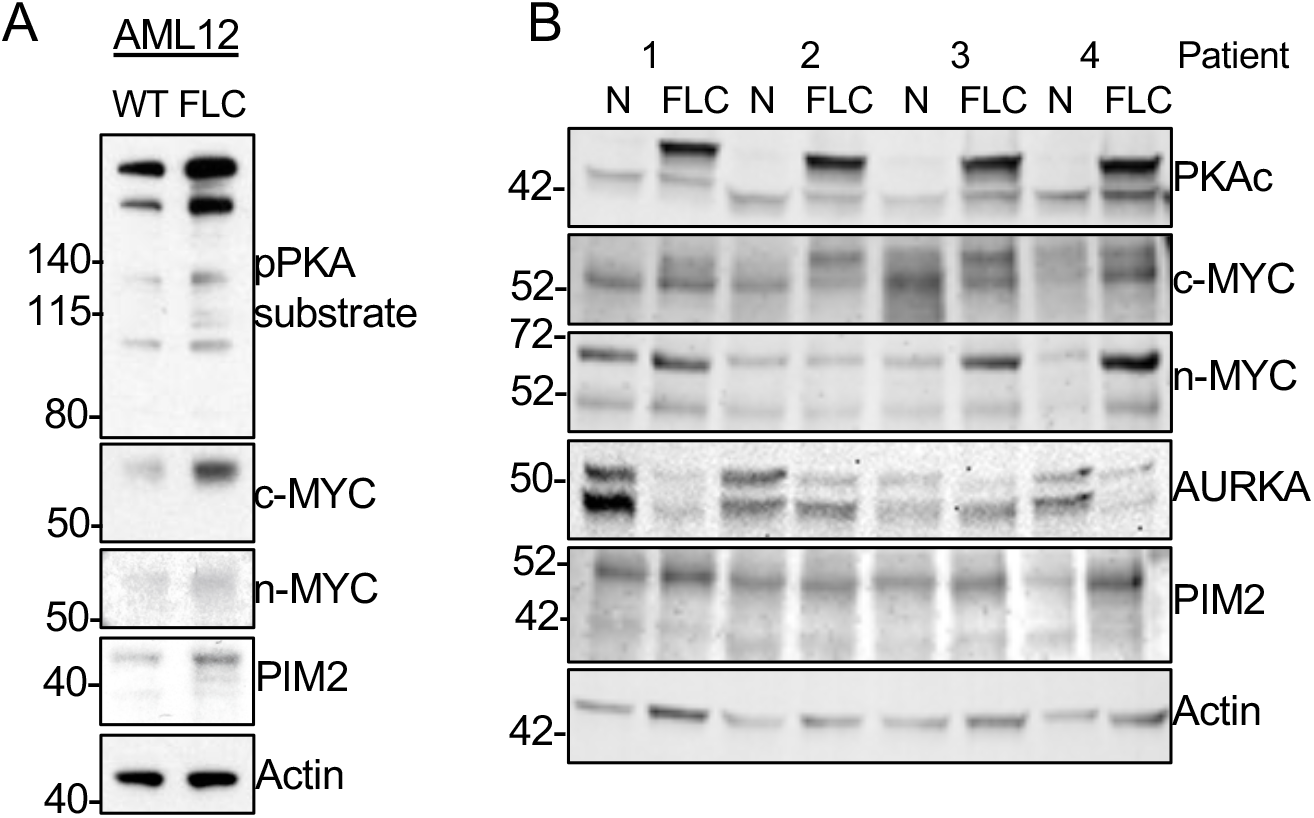
Higher c-MYC and n-MYC protein levels are found in human FLC tumor samples. (A) Immunoblots showing the basal level of PKA activity in phosphorylated PKA substrate and c-MYC, n-MYC, and PIM2 in AML12 WT (left) and AML12^DNAJ-PKAc^ cells (right). (B) Immunoblot showing the presence of DNAJ-PKAc and different level of c-MYC, n-MYC, AURKA and PIM2 in FLC tumor samples (FLC) versus adjacent liver (N) from 4 FLC patients.

Finally, we assessed MYC protein levels in human FLC resection specimens. Here, when compared to adjacent non-transformed liver, we detected not only the DNAJ-PKAc fusion but also increased protein levels of both c-MYC and n-MYC, consistent with our findings in FLX1 cells. However, levels of AURKA and PIM2 were highly variable between tumor and adjacent liver, raising the possibility of additional mechanisms of c-MYC/n-MYC stabilization *in vivo* (Fig. 6B). Thus, while multiple factors may influence c-MYC and n-MYC expression in FLC in vivo, increased expression of one or both is a recurrent feature of this cancer.

## Discussion

Over the last decade, tumor sequencing and mouse modeling studies have demonstrated the importance of GNAS/PKA signaling in cancer, revealing frequent oncogenic mutations in *GNAS* (O’Hayre et al., 2013). Related studies have delineated the essential role of PKAc as its effector (Coles et al., 2020; Patra et al., 2018). Here, we define the context of genetic alterations in *PRKACA* and *PRKAR1A* that result in PKAc activation in cancer and a set of conserved pathways downstream oncogenic PKAc signaling.

Our proteomic analysis has uncovered both expected and novel regulatory effects of PKA modulation in cancer cell lines. In addition to activation of the AKT and BRAF pathways, we noted inhibitory effects of PKA on various kinases, including STK11 and its effectors. We also noted strong modulation of kinases involved in cell migration (eg. YES, EPHB4, LIMK1, LIMK2, and ROCK1). These interesting observations merit further investigation into mechanistic underpinnings of PKA-associated malignancies. Network propagation allowed us to identify two functional clusters in PKA-driven signaling, one driven primary by the RAS/MAPK pathway, and the other by nucleated AURKA, the latter of which was particularly intriguing. AURKA activation is an important step in meiosis, an event that is triggered by PKAc (Frank-Vaillant et al., 2000; Pascreau et al., 2005; Solc et al., 2010). AURKA upregulation has also been noted in GNAS (Coles et al., 2020) and PKA-driven malignancies (Simon et al., 2015). Here, we connect the PKA-AURKA to stabilization of MYC family members, which we demonstrate as a key element of PKA-driven oncogenic proliferation. Of note, prior work on PKA in cancer, particularly in the context of GNAS, has typically been done in systems with concomitant RAS/MAPK pathway alterations (Patra et al., 2018). As FLC rarely bears additional oncogenic mutations (Cornella et al., 2015; Honeyman et al., 2014), the overlapping results from these genetically distinct cell lines suggest that PKAc itself exerts a specific influence in carcinogenesis.

This work highlights the utility of our newly developed FLX1 cells, the first FLC derived cell line, as well as our dox-controlled PKA regulatable system to generate PKA kinome profiles. FLX1 cells were derived from a well-characterized FLC PDX (Dinh et al., 2017), and contain the *DNAJ1-PRKACA* fusion as their primary oncogenic driver. This later feature permitted us to interrogate the direct effects of the fusion kinase. By genetically modulating PKA signaling either by hyper-activation or expression of dominant-negative constructs, we are able to define downstream effectors of PKA activating or inhibiting compounds. Another valuable feature of this approach is the ability to pinpoint PKA independent off-target effects of commonly used PKA-modulating tool compounds. Despite these positive attributes, our models have limitations. For example, PKA signaling is well known to be precisely spatiotemporally regulated (Bauman et al., 2006; Coghlan et al., 1995; Hausken et al., 1996), and our overexpression systems do not allow compartmentalized control of PKA signaling. Additionally, most cell lines used in this study contain oncogenic mutations in addition to their PKA alteration, and it is anticipated that these co-existing mutations may interact with modulated PKA activity to alter the final signaling outcomes. Thus, the identification and development of more PKA-driven cancer cell models is essential.

The data herein support the notion that c-MYC and n-MYC are important effectors of PKA-driven oncogenesis. We found that PKA activity modulates c-MYC protein levels in multiple cell models, and both c-MYC and n-MYC are up-regulated in FLX1 cells. We note that co-expression of c-MYC and n-MYC is relatively atypical in human cancer, with the exception of neuroblastoma (R. Huang et al., 2011), but we found co-induction in human FLC, suggesting that it is characteristic of this cancer. This finding opens a number of interesting avenues for further research. First, while the stabilization of MYC family members by AURKA is well established (Dauch et al., 2016; Gustafson et al., 2014; Richards et al., 2016), mechanistic aspects of this important phenomena are not fully elucidated. PKA activation of AURKA and parallel effects on PIM2 infer that further investigation of the factors that control the relative contributions of these kinases to MYC regulation is a worthwhile endeavor. Similarly, the key transcriptional mediators of the effects of c-MYC and n-MYC on cell growth in FLX1 cells and in FLC remain unknown. Moreover, the extent to which MYC may interact with other key transcription factors such as CRTC2 (Hollstein et al., 2019) or AP-1/CREB (Dinh et al., 2020) unknown. Thus, our discovery of this new means of PKA-responsive MYC activation warrants further investigation.

There is an urgent need for new therapies for PKA-driven malignancies. This challenge is especially pronounced in the case of FLC, where there is no clear standard of care and therapy is not currently tailored to the mechanisms of tumor growth. While further mechanistic investigation and more advanced compounds are needed, our data suggest that CD-AURKAi may improve outcomes for patients with few options, who currently face a grim prognosis.

## Materials and Methods

### Cell culture reagents and treatment

Human bladder 639V cells, human skin Colo741 cells, and human thyroid ML1 cells were maintained in Dulbecco’s modified Eagle’s medium (DMEM) supplemented with 10% fetal bovine serum (FBS), penicillin (100 U/ml) and streptomycin (100 U/ml). The murine hepatocyte AML12 WT and AML12^DNAJ-PKAc^ cell lines were developed as described previously (Turnham et al., 2019). These cells were maintained in 50:50 DMEM/Nutrient Mixture F-12 (F-12) supplemented with 10% FBS, 0.1X ITS liquid media supplement, dexamethasone (0.1 uM) and gentamicin (50 ug/ml). FLX1 cells were derived from a human FLC tumor and xenografted to mice through dispersal and direct plating onto cell culture, and maintained in RPMI with 50 ng/ml HGF, 10% FBS, penicillin (100 U/ml) and streptomycin (100 U/ml). All cells were cultured in a 37°C incubator with 5% CO2. Cells were tested for mycoplasma contamination routinely. Recombinant HGF was obtained from PeproTech; dexamethasone, forskolin, gentamicin, IBMX (3-isobutyl-1-methylxanthine), and 100X ITS liquid media supplement from Millipore Sigma; CD532, DMEM, DMEM/F-12, RPMI, FBS, Lipofectamine RNAiMAX Reagent, Opti-MEM and penicillin-streptomycin from Thermo Fisher Scientific; and MLN8237 and CX-6258 from Selleckchem. The human protein kinase siGENOME siRNA library was obtained from GE Dharmacon. FuGENE 6 transfection reagent and CellTiter-Glo assay system were obtained from Promega. SMARTpool siGENOME siRNA targeting NTC, *DNAJB1, MYC, MYCN, PIM1, PIM2* and *PRKACA* were purchased from Dharmacon.

For individual experiments, cells were seeded at 200,000 cells in 6 cm dishes overnight before treatment, except FLX1 cells, which grew for 2 nights. For drug treatment, a final concentration of 50uM IBMX, 50uM FSK and 1uM of the indicated drug were added to the cells in this order for the desired time periods and harvested. For siRNA treatment, 12 ul of 20 uM siRNA were added to the cells with Lipofectamine RNAiMAX reagent in Opti-MEM, incubated for 72 hours, and harvested.

### Clonogenic Assay

100,000 cells were plated in 6 cm dishes, treated with siRNA for 24 hours, transferred to a 6-well plate at 5,000 cells per well, incubated at 37°C and 5% CO2 for 24 days, and stained with 6.0% glutaraldehyde (vol/vol) and 0.5% crystal violet (wt/vol) in H2O (Franken et al., 2006).

### DNA transfections and lentivirus production

Plasmids containing *PRKACA, PRKAR1A, MYC* and *MYCN* were obtained from the Human ORFeome v8.1 Collection, addgene, or DNASU and cloned into a gateway compatible version of pLVX-Tet-One (puro) with 3xFLAG tag at the N terminal (for *PRKAR1A, MYC* and *MYCN*) or C terminal (for *PRKACA*). The *PRKAR1A*^*G325D*^ single point mutation was introduced using standard PCR technique. The final plasmids were packaged in HEK 293T cells for 72 hours to produce lentivirus, which were used to establish cell lines with respective transgene.

### SDS-PAGE and immunoblotting

Cells were harvested physically and lysed in RIPA buffer with protease and protein phosphatase inhibitors. Protein concentration of cleared lysate was determined by BCA Protein Assay (Pierce). Lysates were separated in 4–12% NuPAGE gradient gels (Thermo Fisher), transferred to nitrocellulose membrane and blocked with 5% milk in TBST using standard technique. Blocked membranes were immunoblotted with antibodies against the following targets separately: AURKA (CST #14475, Biolegend #603301), Phospho-PKA Substrate (CST#9624), c-MYC (CST#18583), n-MYC (CST#84406), PIM1 (CST#3247), PIM2 (CST#4730), PKAC-α (CST#4782), PKAR1a (CST#5675), FLAG (Sigma#F1804), Actin (CST#3700) or COXIV (CST#5247).

Afterward, blotted membranes were washed in TBST, incubated with appropriate HRP-labelled secondary antibodies (CST#7074, 7076), probed with ECL reagents (Thermo Fisher) and developed by x-ray. Blots were washed and stripped with Restore Plus stripping buffer (Thermo Fisher) if multiple probes were required.

### Quantitative RT-PCR

Colo741 and FLX1 cells were seeded and treated in the same manner as described for immunoblotting in preparation for siRNA treatments. RNA was extracted with Trizol reagent (Thermo Fisher) according to the manufacturer’s instructions and quantified with a NanoDrop instrument. Normalized RNA was reverse transcribed with SuperScript II Reverse Transcriptase (Invitrogen). cDNAs were added to PerfeCTa SYBR Green FastMix Reaction Mixes (QuantaBio) and respective primers and analyzed using the BioRad CFX Connect Real-Time PCR Detection System. Primers were designed with Primer3 (Untergasser et al., 2012) and obtained from Elim Biopharmaceuticals (S. Table. 1).

### Cell viability assay

Cells were seeded into 96 well white opaque plates (Greiner) at 2,000 cells per well and incubated at 37°C and 5% CO2 overnight. Cells were treated with selected drugs at different final concentrations and incubated for another 72 hours, though FLX1 cells were incubated for 120 hours due to slow growing speed. After incubation, plates and CellTiter-Glo (CTG) reagent were allowed to equilibrate at room temperature on the bench for 30 minutes. The CTG assay was performed as instructed and measured with the SpectraMax i3 Multi-Mode Platform (Molecular Devices).

### Cell proliferation curves

For experiments with transgenic FLX1 cells, 5333 cells of each line were seeded into black clear bottom 96 well plates (Corning) in 100 ul of media with or without dox (1 ug/ml). After seeding, plates were immediately incubated at 37°C and 5% CO2 inside the Incucyte Zoom system (Essen BioScience) for live cell image and confluence analysis. For experiments with parental cells and siRNA, Colo741 and FLX1 cells were plated and treated with siRNA as described above. Instead of harvesting the cells physically at 72 hours after siRNA treatment, the cells were trypsinized after 24 hours of siRNA treatment and transferred to a black clear bottom 96 well plate at 500 cells per well. The plates were allowed to incubate at 37°C and 5% CO2 for 24 or 36 hours and moved to the Incucyte for further incubation. Once the plates were mounted inside the Incucyte system, pictures of each well were taken every 2 hours for confluence analysis.

### siRNA kinase library screening

384 well plates containing the human protein kinase siGENOME siRNA library (Dharmacon Cat#G-003505) were thawed at room temperature and centrifuged at 1000 rpm for 5 minutes prior to foil removal. 50ul of nuclease-free dH2O was added to each well to reconstitute the siRNA at a final concentration 5uM. Using a Labcyte Echo 525 liquid handling machine, 200nl of reconstituted siRNA from each well from the master plates was transferred to the same position of the corresponding black transparent bottom 384 well daughter plates (Thermo Fisher). Unused aliquoted plates were sealed with foil, covered with plastic lid and stored at -80C. For subsequent experiments, daughter plates with deposited siRNA were thawed at room temperature and centrifuged. 5ul of nuclease-free dH2O was added to each well and agitated at room temperature for 30 min. 10ul of a mixture of RNAiMAX and Opti-MEM was then added to each well and incubated at RT for 20 mins. Finally, 500 FLX1 cells in 30 ul media were added to each well. The plates were transferred to the Incucyte for cell proliferation monitoring.

### TCGA analysis

TCGA PanCancer Project data between 3/13/18 and 4/23/18 were accessed through cBioPortal (at https://www.cbioportal.org) and queried by gene (e.g. *PRKACA, PRKAR1A*). Data were sorted through *Cancer Types Summary* function and exported to Microsoft Excel and Prism for reorganization and analysis.

### Phosphoproteomics

Transgenic cell lines with doxycycline-controlled 3xFLAG-*PRKACA* or *PRKAR1A*^*G325D*^ constructs were treated with DMSO or doxycycline for 48 hours. Cells were then harvested in PBS, lysed in lysis buffer (8uM urea, 50mM Tris pH 8, 75mM NaCl, 1X protease and phosphatase inhibitors) and sonicated at 20% for 15 sec. Bicinchoninic acid (BCA) protein assay was performed to quantify protein lysates. Samples were reduced with 5mM dithiothreitol (DTT), cooled to room temperature, alkylated with 15mM iodoacetamide, quenched with 15mM DTT, diluted with 50mM Tris pH 8 to <2M urea, and subjected to trypsin digestion at 37C overnight. Samples were acidified with 10% trifluoroacetic acid (TFA). 50mg Seppak cartridges were set up on vacuum, and columns were washed with series of MS-grade acetonitrile (ACN), 70% ACN/0.25% acetic acid (AA), and 0.1% TFA buffers. After letting samples drip through columns, columns were washed with 0.1% TFA and 0.5% AA. Samples were eluted and lyophilized in a speed vacuum concentrator, and phosphopeptide enrichment was performed with immobilized metal affinity chromatography following established protocols (Budzik et al., 2020). Phosphopeptides were eluted in 50% ACN/0.1% formic acid (FA) and dried on a speed vacuum concentrator. Enriched samples were analyzed on a Q Exactive Orbitrap Plus mass spectrometry system (Thermo Fisher Scientific) with an Easy nLC 1200 ultra-high pressure liquid chromatography system (Thermo Fisher Scientific) interfaced via a Nanospray Flex nanoelectrospray source. Samples were injected on a C18 reverse phase column (25cm x 75uM packed with ReprosilPur C18 AQ 1.9 uM particles). Mobile phase A consisted of 0.1% FA and mobile phase B consisted of 80% ACN/0.1% FA. Peptides were separated by an organic gradient from 2% to 18% mobile phase B over 94 minutes followed by an increase to 34% B over 40 minutes, then held at 90% B for 6 minutes at a flow rate of 300 nL/minute. MS1 data was acquired with a 3e6 AGC target, maximum injection time of 100ms, and 70K resolution. MS2 data was for the 15 most abundant precursors using automatic dynamic exclusion, a normalized collision energy of 27, 1e5 AGC, a maximum injection time of 120ms, and a 17.5K resolution. All mass spectrometry was performed at the Thermo Fisher Scientific Proteomics Facility for Disease Target Discovery at UCSF and the J. David Gladstone Institutes.

Mass spectrometry data was assigned to human sequences and peptide identification and label-free quantification were performed with MaxQuant (version 1.5.5.1) (Tyanova et al., 2016). Data were searched against the UniProt human protein database (downloaded 2017). Trypsin/P was selected allowing up to 2 two missed cleavages, and variable modification was allowed for methionine oxidation, N-terminal protein acetylation, and phosphorylation of serine, threonine, and tyrosine, in addition to a fixed modification for carbamidomethyl cysteine. The other MaxQuant settings were left as default. Statistical analysis was performed using R (version 3.6.3), RStudio, and the MSstats Bioconductor package (Choi et al., 2014). Contaminants and decoy hits were removed, and samples were normalized across fractions by equalizing the median log2-transformed MS1 intensity distributions. Log2(fold change) for protein phosphorylation sites were calculated, along with p-values. Phosphoproteomic data was uploaded to the PhosFate profiler tool (Phosfate.com, Ochoa et al., 2016) to infer kinase activity. The mass spectrometry RAW mass spectrum files will be deposited into ProteomeXchange via the PRIDE partner repository.

### MIBs

MIBs were performed as described previously (Donnella et al., 2018; Sos et al., 2014). Kinase inhibitor compounds were purchased or synthesized and coupled to sepharose beads using 1-Ethyl-3-(3-dimethylaminopropyl)carbodiimide chemistry. Transgenic cell lines with doxycycline-controlled Tet-on 3xFLAG-*PRKACA* or *PRKAR1A*^*G325D*^ constructs were treated with DMSO or doxycycline for 48 hours then collected in PBS. Samples were lysed in 150mM NaCl buffer with protease and phosphatase inhibitors. Lysates were diluted with 5M NaCl and high-salt binding buffer (50mM Hepes pH 7.5, 1M NaCl, 0.5% Triton X-100, 1mM EDTA, 1mM EGTA). Pre-washed columns containing ECH sepharose 4B and EAH sepharose 4B beads were layered with kinase inhibitor-coupled beads as follows: 200uL JG-4, 100uL VI-16832, 75uL staurosporin, 100uL PP-hydroxyl, 100uL purvalanol B, 50uL GDC, 100uL dasatinib, 50uL sorafenib, 50uL crizotinib, 50uL lapatinib, 50uL SB202190, 50uL bisindolylmaleimide X. Columns were washed with high salt buffer without disturbing bead layers, and affinity purification was performed with gravity chromatography. Bound kinases were washed with high salt buffer, low salt buffer (50mM Hepes pH 7.5, 150mM NaCl, 0.5% Triton X-100, 1mM EDTA, 1mM EGTA), and 0.1% (w/v) SDS in high salt buffer. Samples were eluted twice by capping the column, applying 300uL of elution buffer (0.5% SDS/1% BME/0.1M Tris-HCL pH 6.8) to the column, vortexing, heating to 98C, removing caps, and allowing elution to flow through by gravity. Samples were frozen at -80C overnight, reduced with 500mM DTT, cooled to room temperature, and treated with 500mM iodoacetamide. Methanol/chloroform precipitation, trypsin digestion at 37C overnight, and desalting were performed on all samples. Enriched samples were analyzed on a Q Exactive Orbitrap Plus mass spectrometry system (Thermo Fisher Scientific) with an Easy nLC 1200 ultra-high pressure liquid chromatography system (Thermo Fisher Scientific) interfaced via a Nanospray Flex nanoelectrospray source as described above for global phosphoproteomics. All mass spectrometry was performed at the Thermo Fisher Scientific Proteomics Facility for Disease Target Discovery at UCSF and the J. David Gladstone Institutes.

Peptides were identified with MaxQuant (version 1.5.5.1) and label-free quantification was performed with Skyline (Schilling et al., 2012), with Trypsin [KR|P] selected and allowing up to two missed cleavages. Full scan MS1 filtering was performed with 70,000 resolving power at 400 m/z using the Orbitrap. Statistical analysis was performed with R, RStudio, and MSstats (Choi et al., 2014) to calculate log2(fold change) and p-values of detected kinases. As above, mass spectrometry RAW mass spectrum files will be deposited into ProteomeXchange via PRIDE.

### Network Propagation

The log(2) fold change values of MIBs data and effect size (ES) of Phosfate data for each transgenic cell line treated with or without doxycycline were separately normalized out of one. The union of these two datasets was generated and any duplicate genes were averaged. Z-scores were then calculated and the absolute values of the z-scores for each cell line were separately propagated across the ReactomeFI network using a MATLAB script (J. K. Huang et al., 2018). Propagated heat scores for each gene were multiplied across cell lines containing the same construct (either Tet-on 3xFLAG-*PRKACA* or *PRKAR1A*^*G325D*^), and significance was calculated based on the probability that propagated heat scores match a permuted value by chance. Significant genes (p-value < 0.05) brought out by the network were then extracted and imported into Cytoscape (Shannon et al., 2003). To integrate transgenic cell lines with Tet-on 3xFLAG-*PRKACA* or *PRKAR1A*^*G325D*^, overlapping direct neighbors of *PRKACA* and their interconnections were extracted. The signs of the averaged z-scores of the Tet-on 3xFLAG-*PRKAR1*^*AG325D*^ lines were flipped and averaged with the averaged z-scores of the Tet-on 3xFLAG-*PRKACA* lines, resulting in a final subnetwork for PKAc. Nodes representing the genes were filled to represent the original z-scores which were averaged across cell lines. Networks were searched on Cytoscape for PKAc and its direct neighbors and any interconnections.

### Human FLC samples

Human FLCs and paired normal livers were obtained from the University of Washington Medical Center and Seattle Children’s Hospital after institutional review board approval (SCH IRB #15277). For prospective fresh tissue collections, informed consent was obtained from the subject and/or parent prior to resection. Fresh/frozen human FLC and paired non-tumor livers were homogenized in RIPA buffer with protease inhibitors using a hand-hold Pro200 homegenizer (ProScientific). Protein concentration of cleared lysate was determined by BCA Protein Assay (Pierce). Lysate were separated by 10% TGX gels (Biorad), transferred to nitrocellulose membrane and blocked with 5% milk in TBST using standard technique. Blocked membranes were immunoblotted with antibodies against following targets separately: PKAC-α (CST#4782), c-MYC (CST#18583), n-MYC (CST#84406), PIM2 (CST#4730), AURKA (Biolegend#603301) or Actin (Sigma#A5441). Afterward, blotted membranes were washed in TBST, incubated with appropriate HRP-labelled secondary antibodies (GE Healthcare Life Sciences), washed as before and developed using ECL (Thermo Fisher) on an iBright FL1000.

## Figure legends

**S. Figure 1.**
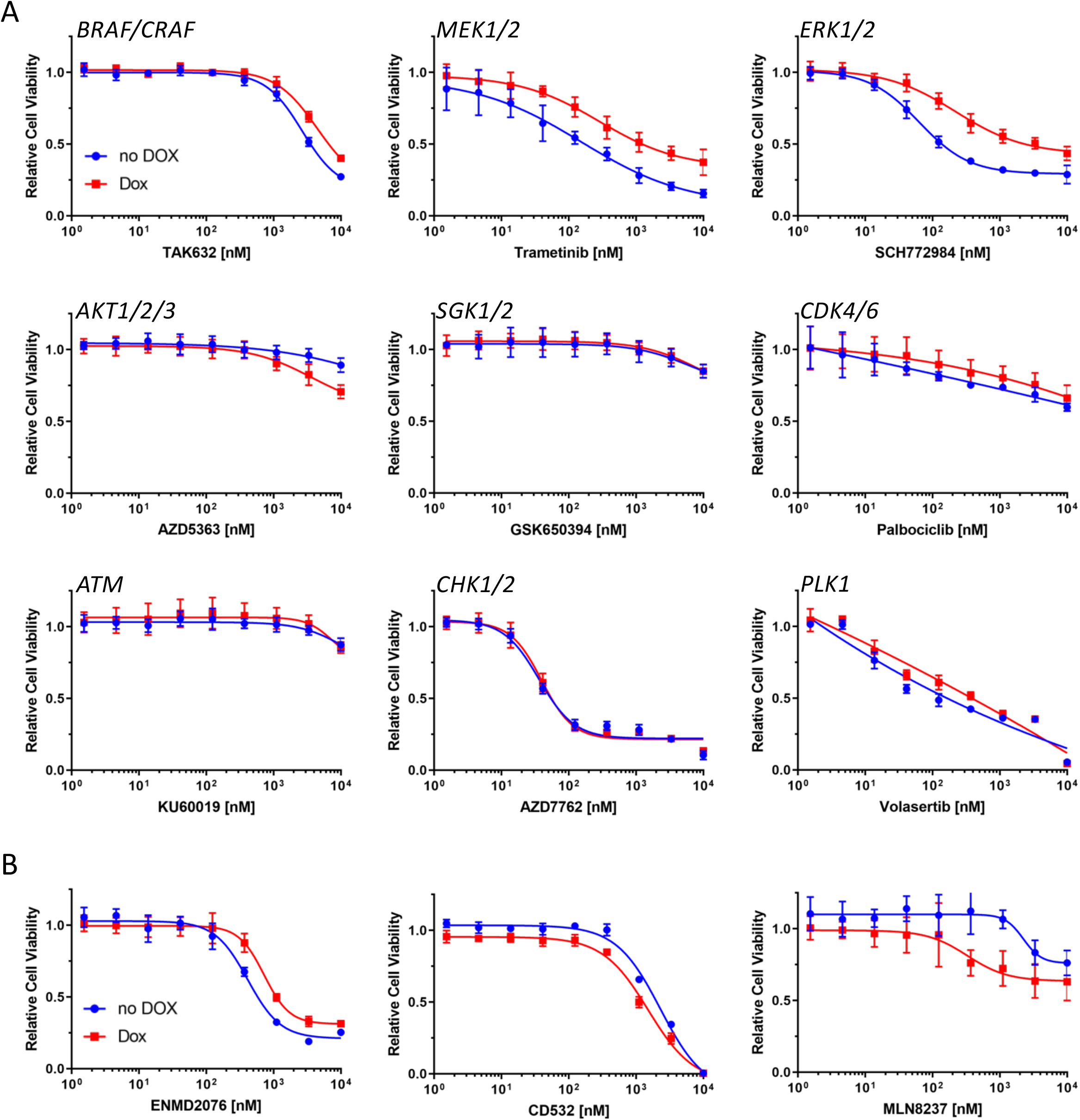
Elevated PKA activity alters drug sensitivity. (A) ML1 cells with Tet-on 3xFLAG-*PRKACA* in 96-well plates with or without dox were treated with various types of inhibitors for 72h at different concentrations, and their relative cell viability was measured by CTG assay versus untreated control samples. Results are the mean ±s.d of one to three biological replicates, three technical replicates per biological replicate. (B) Relative cell viability of ML1 cells with Tet-on 3xFLAG-*PRKACA* with or without dox after treating with various types of AURKA inhibitors for 72h at different concentrations. Results are the mean ±s.d of one biological replicates, three technical replicates per biological replicate.

**S. Table 1.**
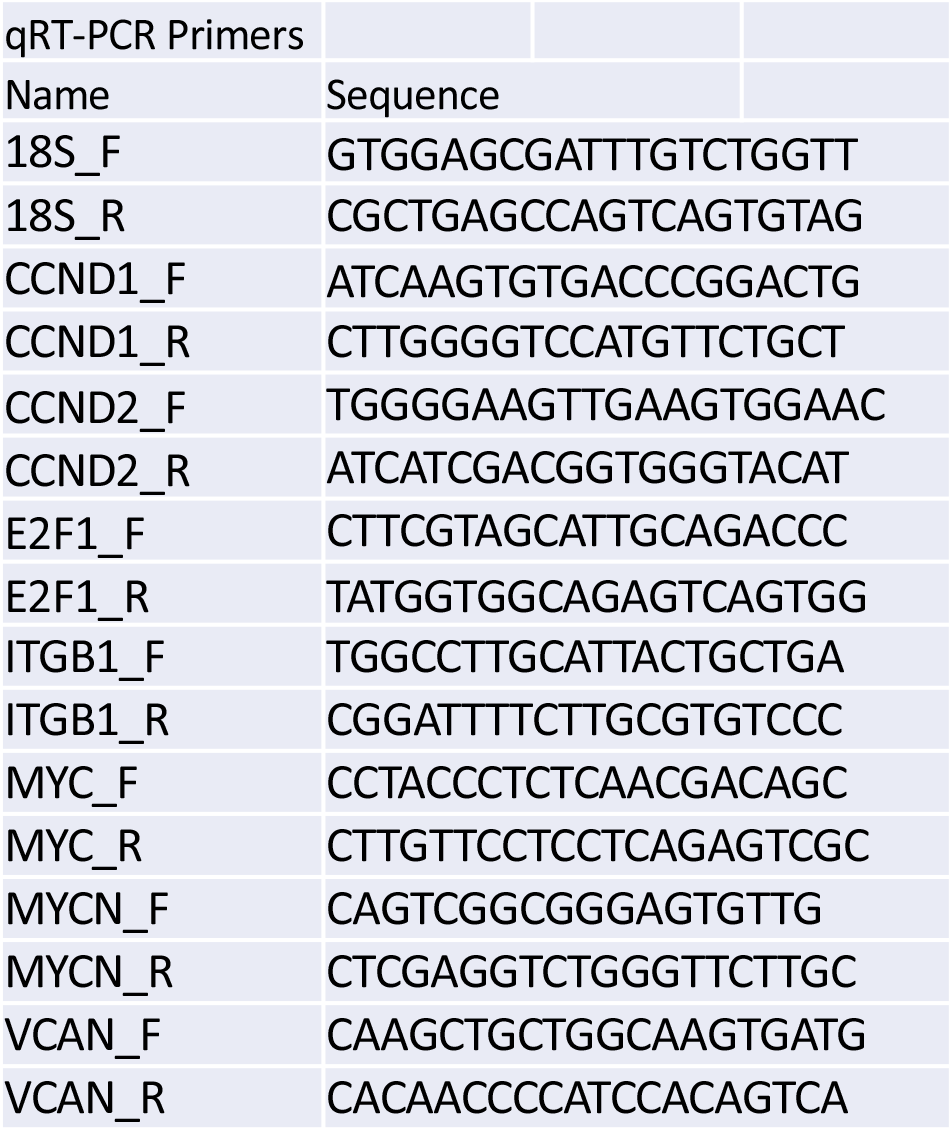
List of primers for qRT-PCR.

**S. Figure 2.**
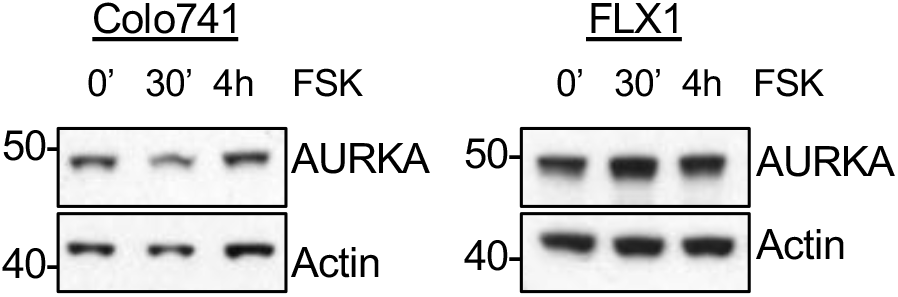
PKA activity has little effect on AURKA protein level. Immunoblots showing the AURKA expression in Colo741 and FLX1 cells after treating with 50uM FSK for 0, 30 minutes, and 4 hours.

